# A Framework for Predicting Design Failures in Engineered Genetic Codes

**DOI:** 10.1101/363812

**Authors:** Bea Yu, Matthew Murphy, Peter A. Carr

## Abstract

Extreme engineering of an organism’s genetic code could impart true genetic incompatibility, even blocking effects of horizontal gene transfer and viral infection. Recent experiments exploring this possibility demonstrate that such radical genome engineering achievements are plausible. However, it is unclear when the modifications will compromise the fitness of an organism. Efforts to reformat an entire genome are difficult and expensive; computational methods predicting fruitful experimental trajectories could play a pivotal role in advancing such efforts. We present a framework for building *in silico* models to assist genome-scale engineering. Genetic code engineering requires choosing from many possible codon-usage schemes, to find a design that is viable and effective. We use machine learning to identify which alternative codon-usage schemes are likely to result in no observed viable cells. Our data-driven approach employs observations of how modifying codon usage in individual genes impacted observed viability in E. coli, revealing salient features for early identification of problematic genetic code designs. We achieved an average area under the receiver operating characteristic of 0.72 on out-of-sample data.

**Author Summary:** As machine learning and artificial intelligence play an increasingly central role in science and engineering, it will be important to establish standardized techniques that facilitate the dialogue between experimentation and modeling. Biological experimental techniques are concurrently evolving at a rapid pace, providing unique opportunities to collect high-quality, novel information that was previously unobtainable. This work navigates the landscape of this vast, new territory, identifies interesting landmarks for exploration and posits new approaches towards advancing our research efforts in these areas. In this work, we show that, using a small dataset of 47 observations and rigorous nested cross validation techniques, we can build a model that makes better-than-random predictions of how codon usage changes in essential genes influence viability in *E. coli.* These predictions can be used to inform experimental trajectories in both genetic code and codon optimization experiments. We discuss ways to improve this model, iteratively, by performing high value experiments that decrease uncertainty in predictions and extrapolation error. Finally, we present novel visualization methods to aid in developing intuitions for how re-coding impacts groups of genes. These methods are also useful tools in building important insights into how well machine learning algorithms can generalize to new data.

## Introduction

With the advent of new *in vivo* DNA editing techniques, engineering at the genome scale is not only achievable but an increasingly inevitable trajectory. In particular, the emerging discipline of genetic code engineering promises novel and far reaching applications in genetic containment, synthesis of proteins incorporating novel amino acids, and immunity to viral infection. Genetic code engineers seek to implement genome-scale changes algorithmically by modifying the rules underlying the genetic code (recoding) and implementing non-natural codes in particular organisms. These changes ultimately result in fundamentally altering how certain codons are recognized during translation. Such “global” changes offer unprecedented opportunities to explore the effect genetic information has on an organism’s specific phenotypes. The first organism recoded genome-wide, C321.ΔA (nicknamed r*E. coli 1.0*), has been engineered by completely removing one of the 64 codons from the genetic code of *E. coli* MG1655 [1,2]. The de-assigned codon has shown facility for being re-assigned to non-standard amino acids [2], for improving resistance to infection by bacteriophage [2] and for providing improved intrinsic biocontainment for engineered microbes [3,4].

Notably, changes to codon usage throughout the DNA of an entire genome are not sufficient to completely implement the desired new genetic code—the gene(s) for the corresponding translation machinery must also be engineered. In this particular case of genome-wide codon removal, deletion of the gene prfA (encoding the release factor protein RF1) was sufficient.

In general, engineering a genetic code genome-wide requires two major steps: 1) completely altering the usage of one or more of the 64 codons throughout the genome, and 2) altering the associated translation machinery to interpret those codons differently. For engineering *rE. coli 1.0*, step 1 required changing the stop codon UAG in all 321 known instances to UAA, a synonymous stop codon, in order to remove the UAG codon from usage throughout the entire genome. Step 2 required deletion of the gene encoding RF1 (release factor 1), removing the cell’s ability to recognize the UAG codon at all.

To perform a similar process with a sense codon, one might alter all instances of the codon GUC (encoding the amino acid valine) to GUG (also encoding valine) and then delete the genes valV and valW, encoding the tRNAs that recognize the GUC codon. These are both examples of a reductionist genetic, i.e. where codons have been cleanly removed from the genetic code. In such cases, further effort would allow the removed codons to be re-assigned to a different amino acid, which again requires engineering both codon usage and the corresponding translation machinery. Importantly, either step 1 or step 2 may result in a growth defect, or lack of any observed growth, in the organism. Choosing codes that minimize this risk is essential when undertaking the long and difficult process of testing a code genome-wide, *in vivo.*

*Lajoie et al.* [5] experimentally explored the impact of step 1 on single genes in *E. coli.* 42 highly expressed genes were redesigned according to a new genetic code that disallowed the use of 13 codons. Individually, each redesigned gene was produced synthetically, inserted into the cell to replace the wild-type version, and doubling time was measured. These redesigned gene sequences also featured substitutions of one allowed synonymous codon for another. For example, instances of the forbidden AUA Isoleucine codon could be replaced with either of the allowed synonyms AUC or AUU, while AUC would be replaced by the only other remaining allowed synonym AUU, because AUA is forbidden. If no colony formation was observed with the altered genes, the impact of changing codon usage in the gene was tested further *in vivo* using alternative designs that departed less drastically from wild-type codon usages.

The experiments in [5] provide an important window into the effects on viability of changing codon usage in highly expressed genes. Other notable efforts by Ostorov et al. in [6] are pursuing the removal of a different, smaller subset of codons throughout the entire *E. coli* genome. This resulting strain, called *rE. coli-57*, would use 57 of the 64 canonical codons genome-wide and may demonstrate increased genetic containment and viral resistance relative to *rE. coli*-*1.0*, in which just one codon was removed genome-wide. Yet the degree of extensive genome engineering required to fully confer genetic containment and viral immunity remains unknown. Identifying the code(s) meeting these objectives—and that can be feasibly engineered while maintaining cell viability—is an extremely complex challenge. The number of codes to test seems unapproachably vast if one considers all possible permutations of codons for each amino acid. Furthermore, modeling the complex mapping of genotype to phenotype is currently limited by incomplete information. Even powerful technologies such as DNA synthesis, MAGE, and CRISPR enabling genetic-code engineering experiments are unlikely to reduce costs in the foreseeable future to the point that all genetic code designs of interest can be evaluated in the laboratory. Under these circumstances, computational methods can play in an important role in establishing an ongoing dialogue between experiment and theory. Results from *in silico* predictive models can be used to identify promising and feasible *in vitro* and *in vivo* experiments. Results from experiments can then be leveraged to increase fidelity and to test accuracy and precision of *in silico* simulations, cf. Figure 1.

**Figure 1.**
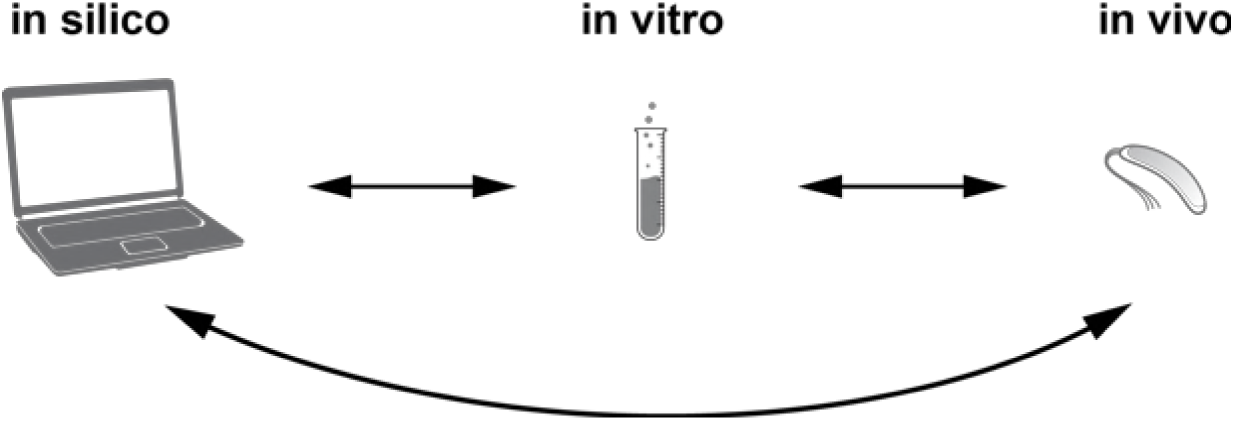
The envisioned tight coupling of information flows among *in silico*, *in vitro* and *in vivo* experiments.

**Figure 2.**
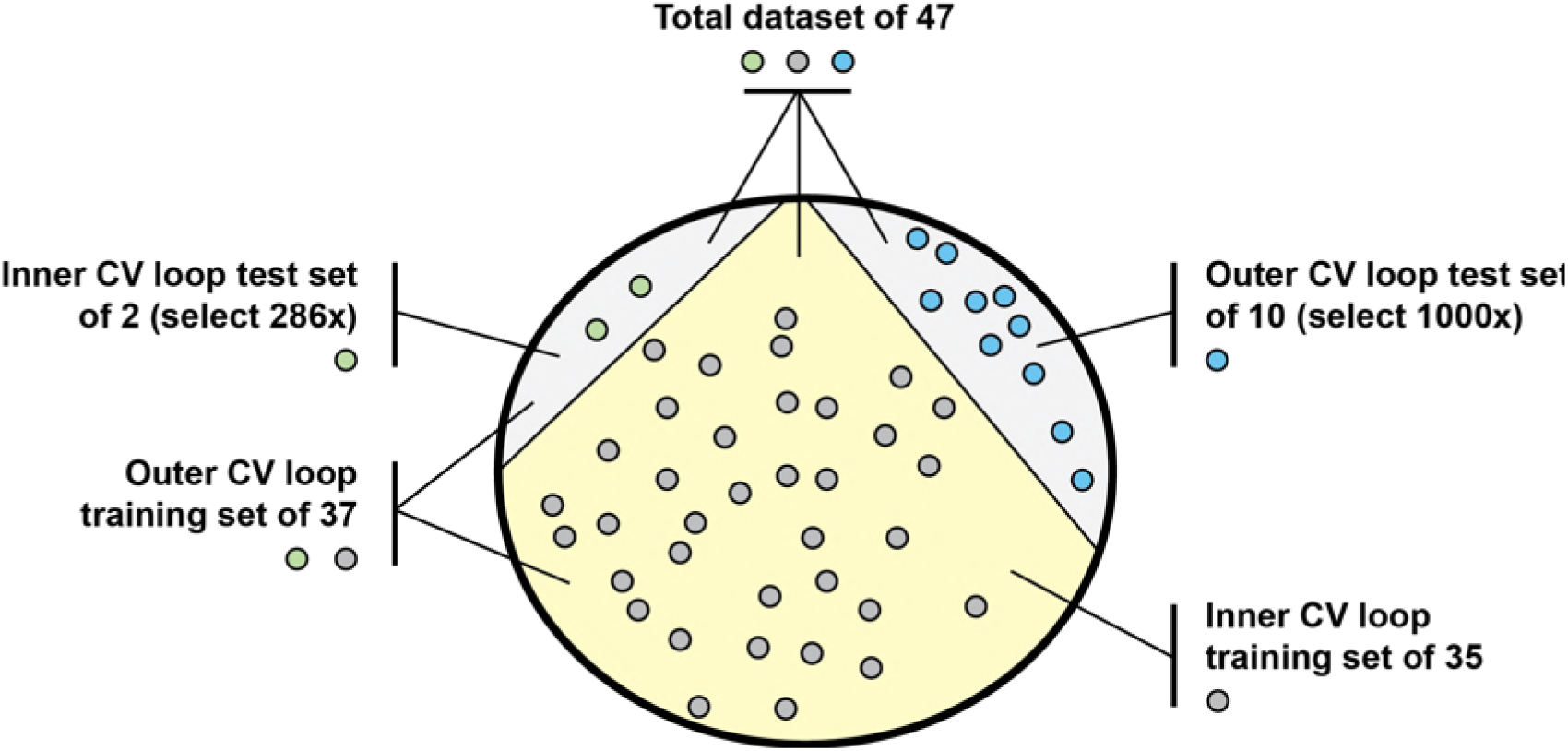
Depiction of data partitions for the nested cross-validation developed for the Gaussian and SVM classifiers. Two partitioning steps occur for each of the 1000 cross validation trials. The first one randomly selects a set of ten hold-out observations (five with no observed colonies, five with observed colonies) from the total set of 47 (blue dots in the diagram), without replacement. Next, from the remaining 37 data points, two are randomly selected, also without replacement, as an inner test sample (one with no observed colony, one with an observed colony) for exhaustive leave-pair-out cross-validation (LPOCV), depicted as green dots in the diagram. The other 35 (gray dots) are used for training the models in the LPOCV loop. This leave-pair-out cross validation is done to learn the optimal number of features allowed in the feature selection process {4, 5, 6, 7}, a classification threshold and whether to include correlated features. The number of features that produces the best AUC from the inner LPOCV is the number of features selected for training a model with the outer CV training set. For details about the nested CV pipeline, see Figure 3.

**Figure 3.**
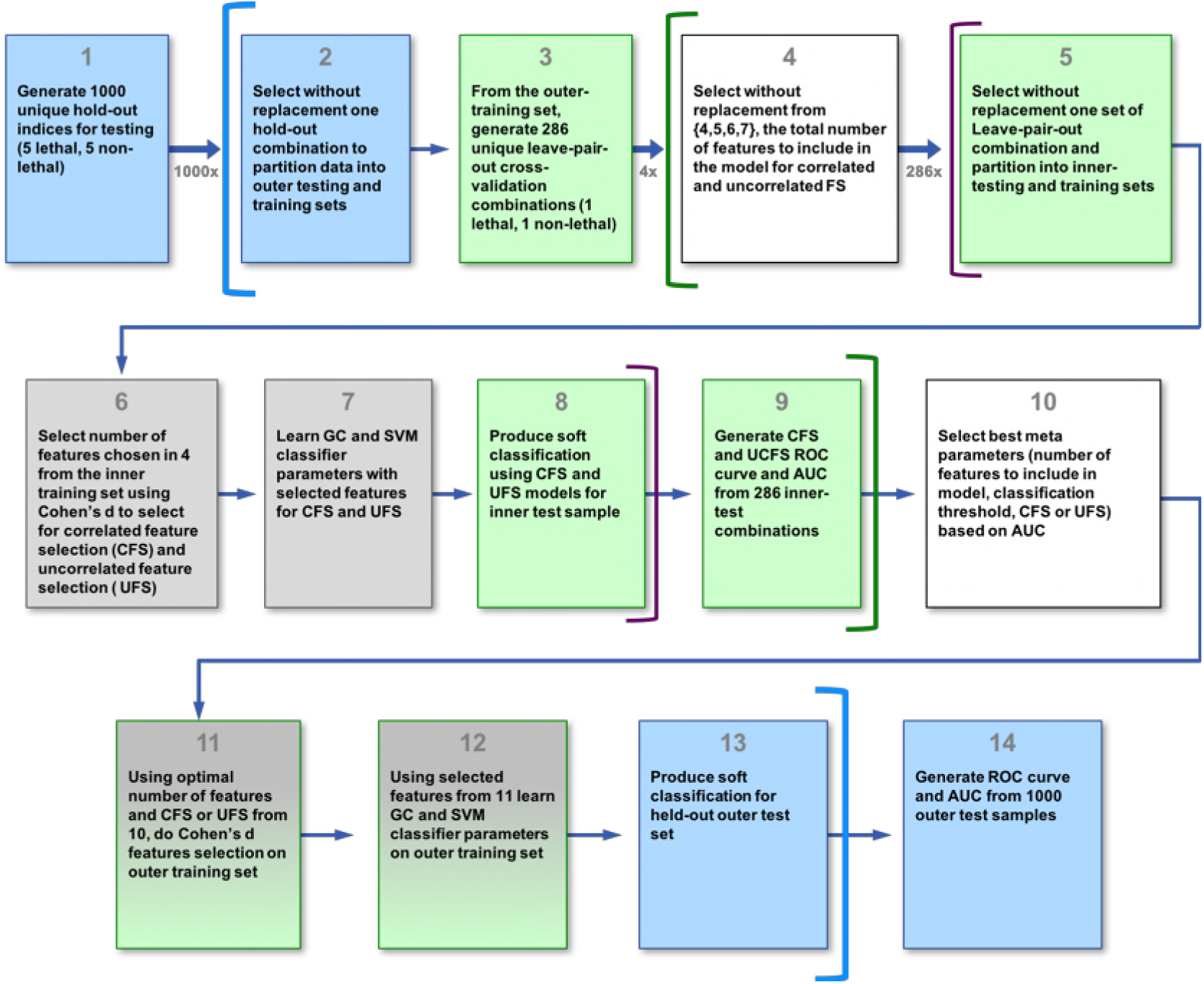
Process diagram of the nested cross validation approach. Block colors correspond to data-type colors in Figure 2 and mean the action described in the block pertains to that subset of data. The inner CV loop chooses the optimal number of features, M, to be used in the outer CV feature selection, which is done on outer CV training data only. Feature selection follows the rubric of choosing the M features with the largest magnitude Cohen’s D effect size for discriminating feature distribution associated with observed and no observed colonies. The inner loop is also used to estimate a classification threshold and whether to include correlated features in the model. These features are then used to train a Gaussian classifier and an SVM. The resulting model is used to predict the class of the held-out ten observations. The outer loop executes 1000 times, resulting in 1000 area-under-the-ROC-curve estimates. To get an estimate of performance, we take the mean of the AUC distribution. KEY: {FS := feature selection, ROC := Receiver operating characteristic, AUC := area under the ROC curve, GC := Gaussian Classifier, SVM := support vector machine}.

*In silico* methods that can predict design failures in step 1 of genetic code engineering would be broadly useful in many genome engineering endeavors. The state of the art in biological computational modeling and analysis, though brimming with promise for explanatory purposes, to our knowledge, lacks the predictive capabilities necessary to address this particular need in genetic code engineering, however. As a baseline, models predicting whether changing codon usage designs in specific genes fail (in the sense that they are associated with no observed growth) in *E. coli* will be particularly useful for saving time and cost entailed in implementing difficult, genome-wide experiments with code designs that are likely compromise its viability. No existing models we know of have this specific capability.

Karr et al. recently published in [7] an *in silico*, whole-cell model of *M. genitallium.* While this type of model may provide useful insights into the effects of changing codon usage in step 1 of recoding, a similar model of *E. coli* is not currently available. Thiele et al. in [8] published a stoichiometric model of *E. coli*’s transcription and translation machinery. This model, however, is not resolved to account for the codon-specific influences in transcription and translation necessary to understand the effects of step 1 on these processes. Lerman et al. in [9] combined the model from [8] with a metabolic model for *E. coli* to capture the influence of genetic information on both metabolism and transcription and translation and, through these processes, phenotype. While this model captures the important influence of metabolism on phenotype, because it uses the transcription and translation model from [8], it lacks the codon-specific resolution necessary to make inferences about phenotypic responses to changes in codon usage entailed by step 1 in the recoding process.

The *in silico* modeling and analysis approaches described above center on digital representations of biochemical and biophysical mechanisms in the cell. An alternative approach involves abstracting the genetic code in terms of its role in information storage and transmission in the cell. With this approach, established physical and mathematical models can be leveraged to explore questions about the driving forces that might have led to the code’s specific topology, defined by its mapping of 61 codons to 20 amino acids. Tulsty et al. model the genetic code as a noisy communication channel and use rate-distortion theory to describe how the code might have evolved to balance the rate and distortion of the channel [10]. Constraints on the code as an information channel, such as “error-load” and amino acid “diversity” (both mapping to the “distortion” of the code “channel”), and the “cost” to the cell of specific binding between codons and anticodons to reduce translation errors (the “rate” of the code “channel”), are likely to influence which genetic code engineering trajectories maintain viability. However, this model does not address factors such as observed organism-specific (e.g. *E. coli* vs. *Plasmodium*) [11] and category-of-gene specific (e.g. highly expressed vs. not highly expressed) [12, 13] codon biases that are impacted when synonymous codons are substituted in step 1 of the recoding process. Modifications in step 1 that alter these biases are also likely to influence the viability of an organism.

To address this capability gap in the state of the art of *in silico* modeling, we developed a phenomenological model specifically to predict the consequences of different codon-usage changes on observed cell growth (one measure of a design success or failure). For the set of individual, highly-expressed, essential genes used in [5], we computed a set of features that address both physio-chemical and information-theoretic characteristics of each gene’s DNA sequence. From these, leveraging accompanying observations of colony formation from [5] in binary form (i.e. {yes, no}), we determined a salient subset of features and learned a statistical model that predicts whether changing codon usage in a gene will impact *E. coli*’s ability to form observable colonies when grown on agar plates. To assess generalizable performance of a model trained with this relatively small dataset, we devised a performance evaluation method using nested cross validation (CV) with random subsampling. The best model yielded an average AUC of 0.72 and a mean sensitivity of 0.80.

We used this model to make design success/failure predictions for *E. coli* genes with codon usage altered *in silico* according to several alternative, novel algorithms. The algorithms we chose incorporate domain knowledge about interesting edge-cases in possible genetic code designs. This model predicted that two out of nine re-coding algorithms are likely to fail upon *in vivo* implementation. The classifier provided information about the effects of changes in codon usage in *individual genes* on cell viability. To gain insights into the effects of algorithmically modifying codon usage in *sets of genes*, we produced scatter-plots of salient features computed for the genes, bounded by convex hulls. This visual representation provides a gestalt view of how different algorithms impact the same group of genes. It is possible with this technique to visually gauge similarity between algorithms in terms of how close their convex hulls are in the feature space and whether these convex hulls are similar in shape and size. With these plots, we also arrived at a qualitative representation of expected extrapolation error from models learned on the Lajoie gene/code dataset in [3].

## Results

### Nested CV results

The average AUC, sensitivity and specificity over the 1000 trials for the GC and SVM classifiers appear in Table 4. Thresholds determining sensitivity and specificity were learned on out of sample data from the inner CV loop. The AUC of the GC classifier, AUC = 0.72, was significantly better than that of the SVM, AUC = 0.68, at the 5% level. Both AUC values were significantly better than random, at the 5% level. The sensitivity of the GC classifier was 0.8; that of the SVM, 0.76. Specificity of the SVM was higher, 0.52, than that of the GC, 0.48. Given the problem space we are addressing with this model, the GC appears to be a better choice for discriminating design success/failure than the SVM because it has a higher true positive rate. We wanted to minimize the false negative rate (1 - true positive rate), e.g. falsely classifying a modification that would result in design failure as a modification the would result in a successful design, because *in vivo* experiments are expensive and only modifications with a high probability of viability are likely to be good candidates for full, *in vivo*, recoding experiments.

### Classifying the effect of the nine new codon-usage algorithms on the 40 genes in vivo

Figure 7 shows split bean plots of the features with the strongest Cohen’s d selected using the uncorrelated feature selection method on the entire FCS and FC dataset, as described above.

Cyan and yellow lines indicate the log of the computed feature values, blue and orange contours are kernel density estimates of the log feature observations, dark blue and light brown solid lines indicate mean log feature values associated with design success and design failure, respectively, and the black dotted line indicates a grand mean over all log feature values plotted. For the SVM LPOCV parameter selection step, we achieved the best predictive performance using five uncorrelated features selected based on the magnitude of their Cohen’s d, as opposed to using all features, including correlated ones. For the GC, five features chosen from all features, including correlated features, performed best in the LPOCV. Three of the five features selected for the GC were the same as those selected for the SVM: the Hamming distance between the base pair distribution of the WT and modified sequence (HammingBPS), the KL divergence between the codon distribution of the WT and modified sequence (KLdivCodons) and the ratio of CAI from the 31^st^ codon to the final codon before the stop codon (ratioCAIend30codonsPlus). The remaining two CAI features selected for the GC were highly correlated with ratioCAIend30codonsPlus, had very similar bean plots and are therefore not shown.

Since the magnitude of the Cohen’s d was used to select features, features with, on average, very different values for the design success/failure classes, and small, pooled standard deviation, were likely to be selected. In Figure 4, it is clear that the large mean difference between the ratioRNAFoldRBS30 of the two classes is offset by the large pooled standard deviation, making the magnitude of Cohen’s d for this feature one of the lowest in the five selected features. This feature, as well as normMICodons, was not selected in the GC correlated variable feature selection routine. In contrast, ratioCAIend30codonsPlus has a large mean difference between the two classes paired with a relatively small, pooled standard deviation, making it the feature with the strongest Cohen’s d of the five. The ratioCAIend30codonsPlus is the codon adaptation index ratio for the part of the gene following what has been called a “codon ramp” [19], the beginning of the gene characterized by, on average, codons that are translated more slowly. It is not clear why the CAI from the latter part of the gene was more discriminatory for design success classifcation in the FC and FCS modified dataset.

**Figure 4.**
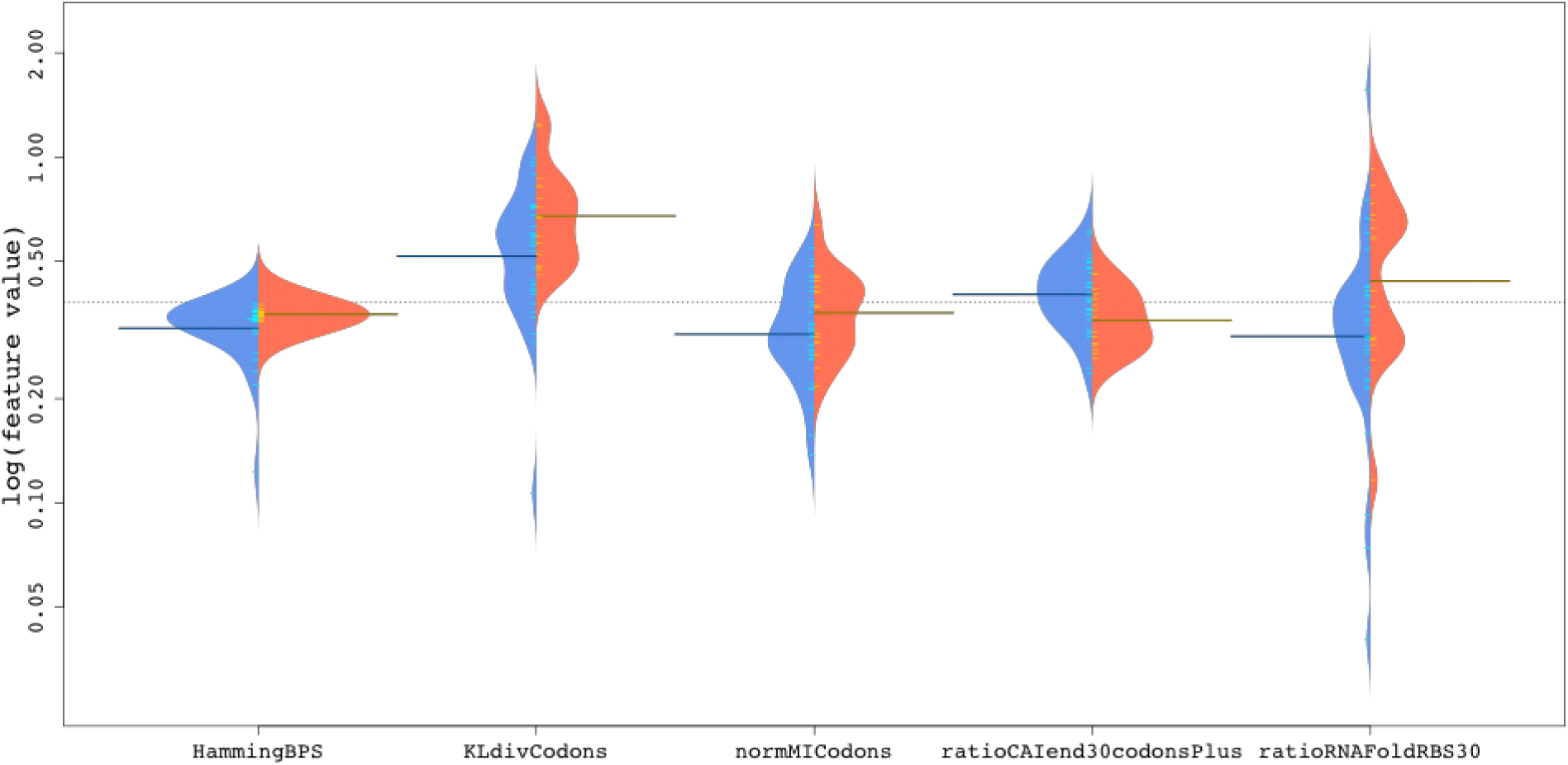
Split bean plots of the five uncorrelated features selected for the SVM classifier. Three of the five, the Hamming distance between the base pair distribution of the WT and modified sequence (HammingBPS), the KL divergence between the codon distribution of the WT and modified sequence (KLdivCodons) and the ratio of CAI from the 31^st^ codon to the final codon before the stop codon (ratioCAIend30codonsPlus), were also selected for the GC correlated feature set. The other two features selected for the GC classifier are highly correlated with the CAIend30codonsPlus, have similar feature histograms and are therefore not shown. The blue envelope is the kernel density estimation of the feature histograms for the modified genes that resulted in a successful design while orange is that of the modified genes that did not. Cyan and yellow lines signify the log of feature values for the two classes of features, respectively. Dark blue and brown long lines indicate mean log values for features associated with design success and failure, respectively.

The Hamming distance for the base pair distribution and the KL divergence for the codon distribution were also selected in both the GC and SVM feature selection methods. The Hamming distance between base-pair distributions measures the minimum number of base pair substitutions that would be required to change the modified sequence back to the wild-type sequence. Modifications that failed had, on average, more base pair substitutions than modifications succeeded. KL divergence can be interpreted as measuring the difference between two sets of observed data by quantifying the amount of extra information required to compress the information in one set using a scheme based on the generating distribution of the other set [31]. It is not a symmetric measure and therefore not a proper distance. The KL divergence between the codon histogram of the wild-type codon frequencies and the codon histogram of the modified gene codon frequencies is therefore the amount of extra information required to compress the codon frequency information from the wild-type sequence using a scheme optimized for the modified sequence. The strong, positive Cohen’s d for this feature in this dataset entails that design failure is correlated with a higher wild-type KL divergence from modified versions of a gene. Thus, the amount of extra information required to compress the codon frequency information from the wild-type sequence using a scheme optimized for the modified sequence was higher on average in modified genes associated with design failure relative to modified genes associated design success.

In Table 5., the designed genes predicted by the SVM and/or the GC to fail are shown. Of the nine algorithms, implementation of two were predicted to fail for one or more modified genes. The Min_weak_ algorithm had the biggest impact on viability-twenty-eight out of the forty genes modified with this algorithm were predicted using either the GC or SVM to fail. tRNA_sub2_ was similarly predicted to fail for eleven out of forty genes. Sub-11 is not predicted to result in a design failure by either classifier for any of the forty genes. The codons removed in *rE. coli*-*57* are a subset of those removed in Sub-11 [6]. It is worth noting therefore that, when implemented individually in the forty genes in this study, the alternative genetic code in [6] would also be predicted by our models to succeed. Ostrov et al. plan to remove seven codons (cf. Table 3) from the entire *E. coli* genome, simultaneously. If there is no observed colony formation in *rE. coli*-*57*, our results suggest that this may to be due to interactions between the recoded genes and/or the individual effects of recoding genes not included in our study.

The GC is more conservative than the SVM in classifying failed modifications for this new set of algorithms. It performed significantly better, using the AUC metric, than the SVM in the nested cross-validation study on the FC + FCS dataset (Table 4). Although the two classifiers agree that many of the Min_weak_ modified genes will fail, there are discrepancies as well. All but two of the modified genes classified as failing under the GC were classified similarly by the SVM. Notably, rplL was the only gene modified with tRNA_sub2_ that was classified as failing by the GC. This gene was not classified similarly by the SVM when modified by any of the algorithms we tested. Though the GC performs significantly better than the SVM in the validation results, the SVM still performs significantly better than random. Therefore, the SVM may provide complimentary information to the GC and the results from the two classifiers could be combined to provide more robust discrimination.

Existing methods could be applied to automatically combine classifier predictions. For example, a majority-rule-like, classifier combination could be defined such that, if the GC and the SVM each classify at least one gene as preventing colonies from forming for any algorithm, that algorithm should be eliminated from the suite of potential algorithms to test *in vivo.* This rule would eliminate both the Min_weak_ and the tRNA_sub2_ algorithms as candidates for *in vivo* genetic code engineering experiments. Such a rule could be honed to weigh the decision rule to throw-out a potential algorithms to test *in vivo* more heavily if multiple genes are classified by each classifier as failing for a given algorithm. Although not shown in Table 4., we tested random forest, gradient boosting and extra trees classifiers from Scikit Learn on this dataset as well. None of these approaches outperformed the SVM and GC classifiers, implemented in Matlab. In principle, however, with further parameter tuning, performance with these techniques might be improved and results combined with the SVM and GC classification outputs to further improve performance.

In the absence of an explicit model combining the predictions of the two classifiers there is enough consensus between them to suggest strongly that tRNA_sub2_ and Min_weak_ should be eliminated as candidate algorithms to be used in genetic code engineering experiments. Although extreme algorithms such as Min_weak_ and tRNA_sub2_ might confer strong viral resistance and genetic isolation to engineered organisms, they don’t perform well in the colony-formation dimension of the multi-objective optimization genetic code engineering attempts to solve. Observed colony formation is a necessary condition that must be met before other dimensions in the problem space defining optimal algorithms, such as viral resistance, genetic isolation and non-natural amino acid and protein generation can be examined.

In Table 5., we also gain insights into which genes are most likely fail in *E. coli* when modified with any algorithm. We call these types of genes “bellwether genes” in reference to the idea that one or a few sensitive genes could be used as a first pass to evaluate algorithms *in vivo* before undertaking more comprehensive, expensive genome-scale modifications. If a modified bellwether gene fails when implemented *in vivo*, the algorithm can be discarded, if not, the algorithm can be evaluated further for suitability. rplB, rpsC, rplO, and rpsA appear to be promising “bellwether gene” candidates since changes to them are notably predicted to fail for tRNA_sub2_, by the SVM, and Min_weak_, by the SVM and GC. These genes are also from the set that resulted in no colony formation *in vivo* when modified with the FCS algorithm. Increasingly comprehensive models trained with more data will enable us to define this set with more confidence.

### Visualizing effects of changing codon usage on sets of genes

In Figure 5, we present an example of a new visualization technique we developed to facilitate gestalt understanding the effects of algorithms used for genetic code engineering on groups of genes. The axes of the plots used in the visualization can be any features determined to be salient in genetic code or genome engineering. For this work, we focus on features useful for design success/failure classification, as measured by observation/no observation of colony growth, discovered using our feature selection routine described above. Two of the five selected features serve as axes in Figure 5. These two features can be plotted for each gene from the set of 40 genes in Table 1. modified with any number, N, of codon-usage algorithms, resulting in N points per gene, 40 points per algorithm and N^*^40 points in all. To reduce complexity as we describe components of the visualization, we show only features from genes modified with two of the nine novel algorithms presented above, Min_weak_ and tRNA_sub1_. The 40 points from each algorithm are bound by line segments defining the convex hull (yellow for Min_weak_ and blue for tRNA_sub1_) in order to visually group the influence of each algorithm on the set of genes in the salient feature space. Generally, the convex hull of a set of points in a Euclidean space is defined as the smallest convex set that contains that set.

**Figure 5.**
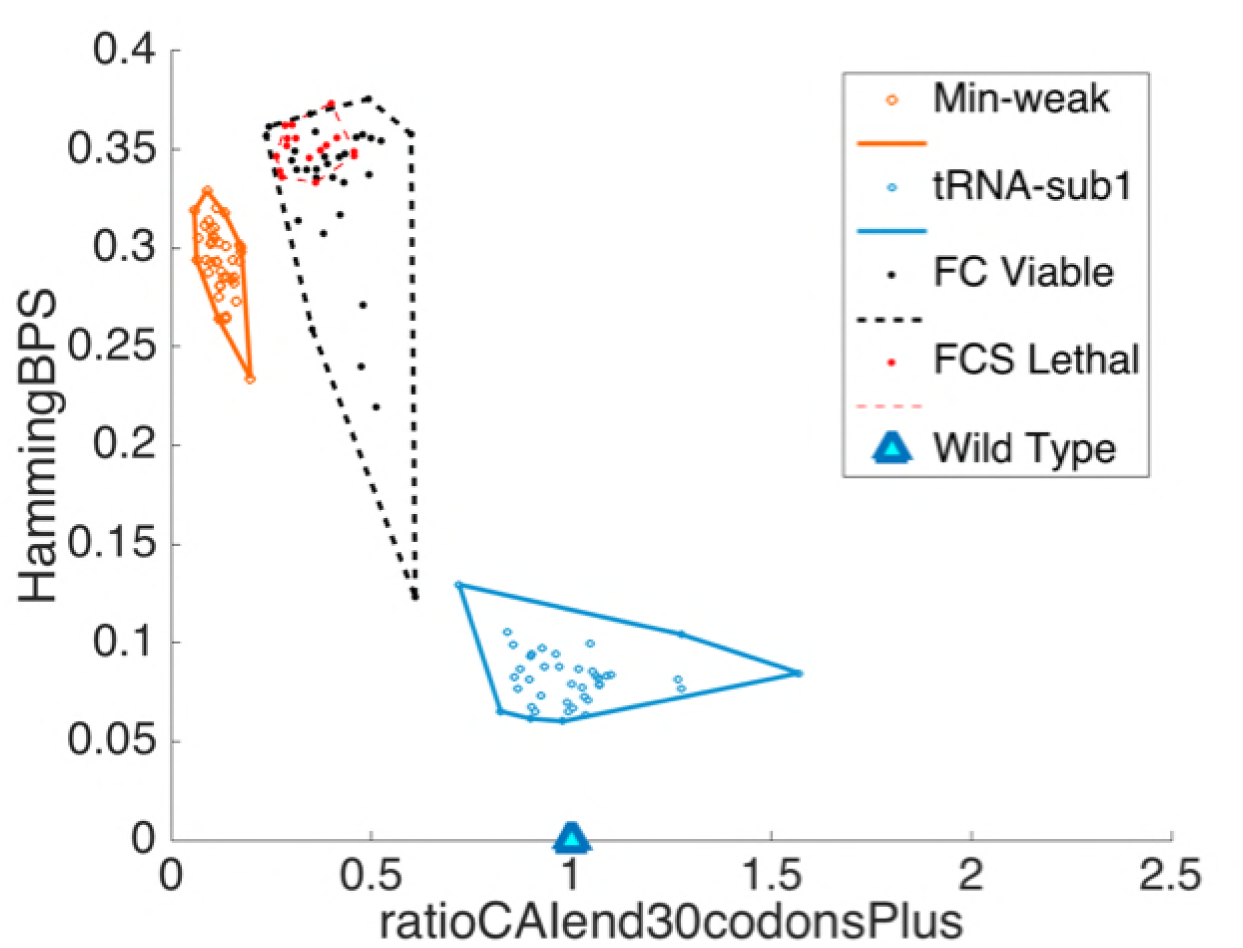
A data visualization method using convex hull plots in salient feature dimensions. Here, we show convex hulls circumscribing the feature values from 40 genes from Table 1, modified with two of the novel algorithms, Min_weak_ and tRNA_sub1_. The features are two of the five selected in the feature selelction routine for design success/failure prediction. This visualization has two benchmarks to orient investigators-the “wild-type origin” and the convex hull plots of features for the the “failed” (red dashed) and “successful” (black dashed) training samples. The wild-type origin shows the feature value a gene would have if it was not modified. The amount the feature of an modified gene deviates from the wildtype origin provides insights into how “different” an modified gene is from the wild-type sequence. The training data convex hull plots show what parts of the feature space are inhabited by observations of successful and failed designs. If genes modified with a new algorithm have features near the training examples that resulted in a design failure, they have the potential to result in a design failure as well.

**Table 1.**
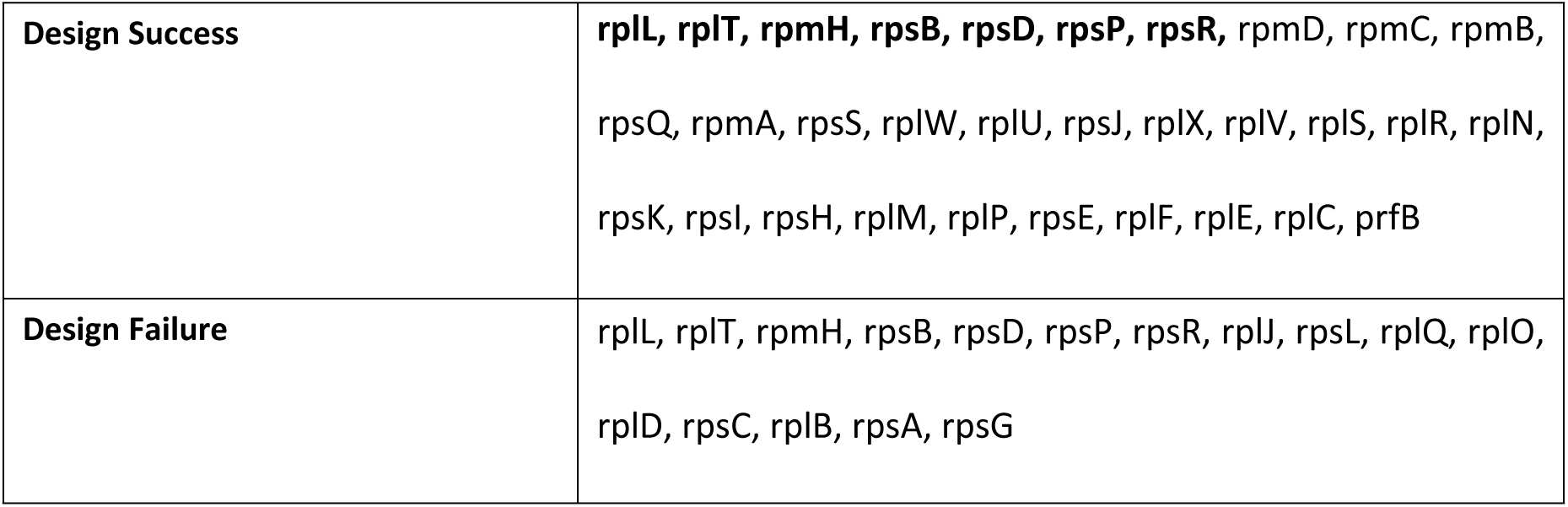
Genes used in this work for model development and performance evaluation and for recoding with nine novel codes and generating predictions and visualizations with sequence data from [3] and EcoCyc. Genes in represented in bold, in the **Design Success** row, resulted in no observed colonies when modified with the FCS algorithm and implemented in *E. coli*, but resulted in observed colonies when modified with the less extreme FC algorithm [3]. These genes also appear in the **Design Failure** row, but in normal type. Normal type designates genes modified with the FCS algorithm, bold, genes modified with the FC algorithm. Genes represented in bold are in both the successful and failed design datasets because they were modified with two different algorithms in [3].

|  |  |
| --- | --- |
| **Design Success** | **rplL, rplT, rpmH, rpsB, rpsD, rpsP, rpsR,** rpmD, rpmC, rpmB,<br>rpsQ, rpmA, rpsS, rplW, rplU, rpsJ, rplX, rplV, rplS, rplR, rplN,<br>rpsK, rpsI, rpsH, rplM, rplP, rpsE, rplF, rplE, rplC, prfB |
| **Design Failure** | rplL, rplT, rpmH, rpsB, rpsD, rpsP, rpsR, rplJ, rpsL, rplQ, rplO,<br>rplD, rpsC, rplB, rpsA, rpsG |

This visualization approach contains several benchmarks to orient investigators to important features of the algorithms. First, a “wild-type” origin is depicted as a cyan triangle in each pairwise plot. This represents the feature values that an unmodified gene would have and can be used to measure how distant algorithms are from wild-type configurations. Because most of the features we used in this work are ratios or normalized distances, the “wild-type origin” for any given feature is 1 or 0. Because all of our features were distances or ratios, the wild-type origin is the same for every gene and can be represented as one point in the pairwise feature space. In Figure 5, the Min_weak_ algorithm, in which 43 codons are removed from the code, moves the set of 40 genes far from the wild-type origin at (1,0), whereas the same set modified with the less extreme tRNA_sub1_ algorithm, in which 28 codons are removed, is relatively much closer to the origin.

Second, relevant groups of the training data are plotted using dashed convex hulls-one in red, to indicate examples resulting in design failure, the other in black, indicating examples of successful designs. These convex hulls demonstrate where in the feature space the model had information about colony formation from which to learn classifier-model parameters. Using this information, areas of the feature space lacking training examples can be identified and strategically populated with data in future experiments specifically designed to make the model more robust and comprehensive. Predictions for new examples far from the training data will likely have more extrapolation error and should be treated with more skepticism.

Third, feature landscapes of how modifications effected colony formation are defined with the dashed convex hulls, allowing investigators to immediately identify algorithms likely to result in a design failure by sight. In Figure 5, the Min_weak_ convex hull is close in the feature space to the red, dashed convex hull bounding the examples that resulted in a design failure in the training set. This was the novel algorithm predicted to fail in the greatest number of genes from the set of 40. On the other hand, the tRNA_sub1_ algorithm convex hull lies far from that bounding the training examples associated with no observed colonies.

Finally, the relative placement and size of the convex hulls of different algorithms are also informative. In Figure 5, the convex hull defining the features from the Min_weak_ algorithm is quite far from that defining features from tRNA_sub1_, consistent with the fact that the two are different “types” of algorithms in the sense that they change the features of the same set of genes differently. Algorithms can also be distinguished by the sizes of their convex hulls. That of the Min_weak_ algorithms is small compared to the convex hulls bounding the training data associated with observed colonies and the tRNA_sub1_ algorithm. This means that, even though the Min_weak_ algorithm is extreme in the number of codons it excludes, it changes all 40 genes away from wild-type configurations “in the same way” in the salient feature space. tRNA_sub2_, though less extreme, influences the genes in the set “in different ways” in the feature space, leading to more dispersion and a larger convex hull. In Figure 6a-c, we plot the convex hulls of all 9 algorithms applied to the set of 40 genes from Table 1. As would be expected, the convex hulls of the features of the forty genes recoded with the Min_weak_ algorithm lies closest to the red, dashed convex hull bounding the examples from the FC algorithm that resulted in design failure in all three pairwise feature plots. Also, as would be expected, the convex hull bounding the gene set recoded with tRNA_sub2_ inhabits parts of the feature space close to those inhabited by the genes recoded with Min_weak_. The convex hulls of these algorithms are plotted with thicker lines to indicate that they are predicted to fail if implemented in one or more of the genes. In particular, both of these algorithms result in a ratioCAIend30Plus less than 1 for all 40 genes, and less than 0.5 for most of the genes, indicating the CAIend30Plus for the modified genes is much reduced relative to that of the wild-type genes. Since the reference set for the CAI consists of highly expressed genes, the higher the CAI, the more likely it is that a gene is highly expressed. This set of 40 genes was selected based on the fact that they were essential and highly expressed. Therefore, the Min_weak_ and tRNA_sub2_ changed the codon composition of these highly-expressed genes in the wild-type organism so that their codon composition was more similar to that of poorly expressed genes, a change that would be intuitively be expected to fail. Furthermore, it is worth noting that the ratioCAIend30Plus had a higher Cohen’s d than the ratioCAI, computed over the entire gene. While more data are needed to confirm this with significance, these results suggest that eliminating the “codon ramp” from the CAI computation may lead to a more sensitive metric for predicting design success.

**Figures 6 a-c.**
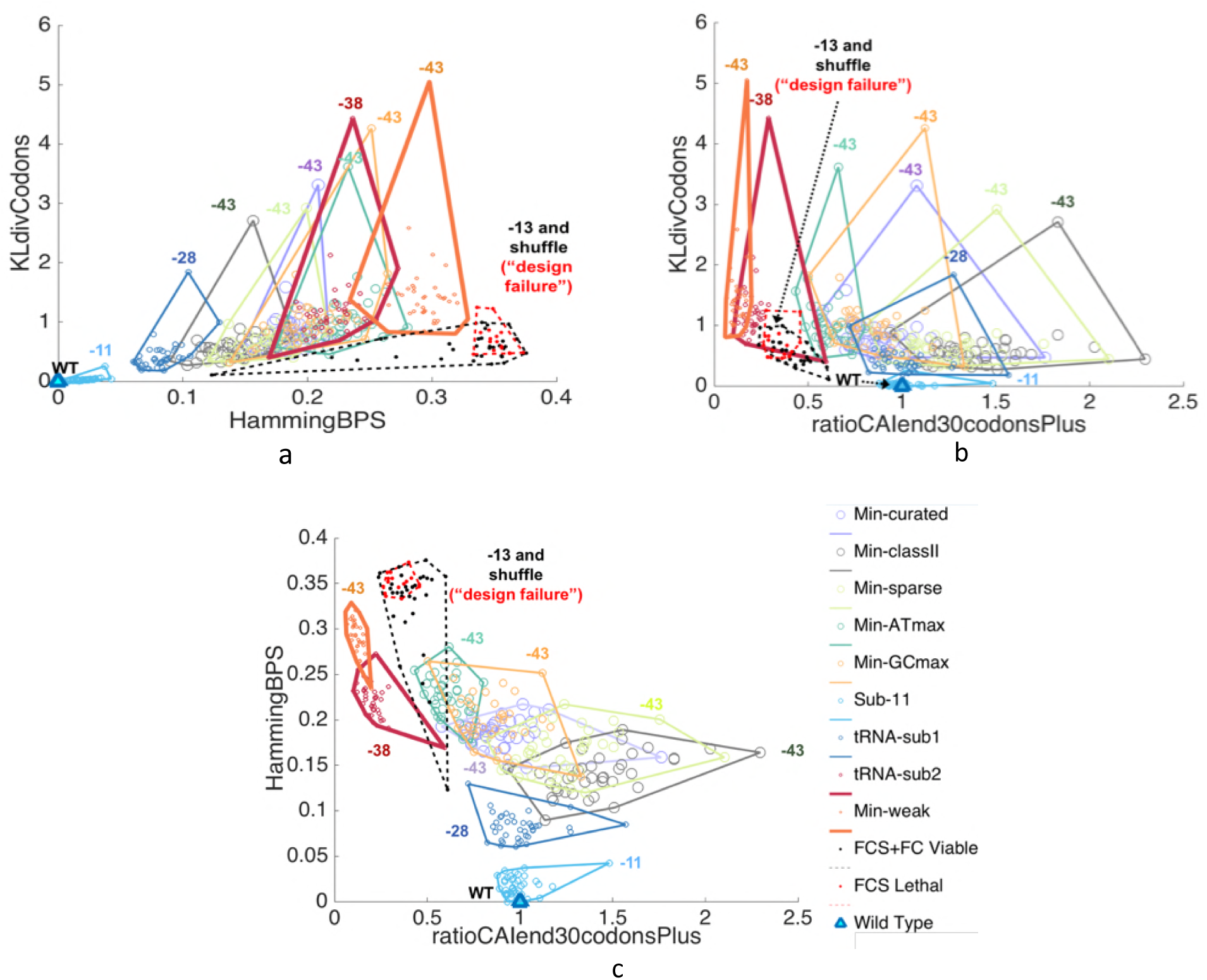
Pairwise convex hull plots in three feature dimensions selected in both the GC and SVM feature selection routines. Each convex hull circumscribes the same 40 genes but modified with each of the nine, novel algorithms. Note that the two algorithms predicted to fail when implemented in particular genes (bolded convex hulls), Min_weak_ and tRNA_sub2_, have similar feature values for this set of genes. Also note that the more extreme the algorithm, the more features deviate from WT features (cyan triangles). In general, predictions for unseen data far from the data on which the model was learned (in the dashed convex hulls), should be treated with more caution than those for unobserved data in the domain of the training data. Note in 9a that rpsG is an extreme outlier for all algorithms in the KLdivCodon dimension (point in each algorithm’s convex hull with largest KLdivCodon).

**Figures 7 a-c.**
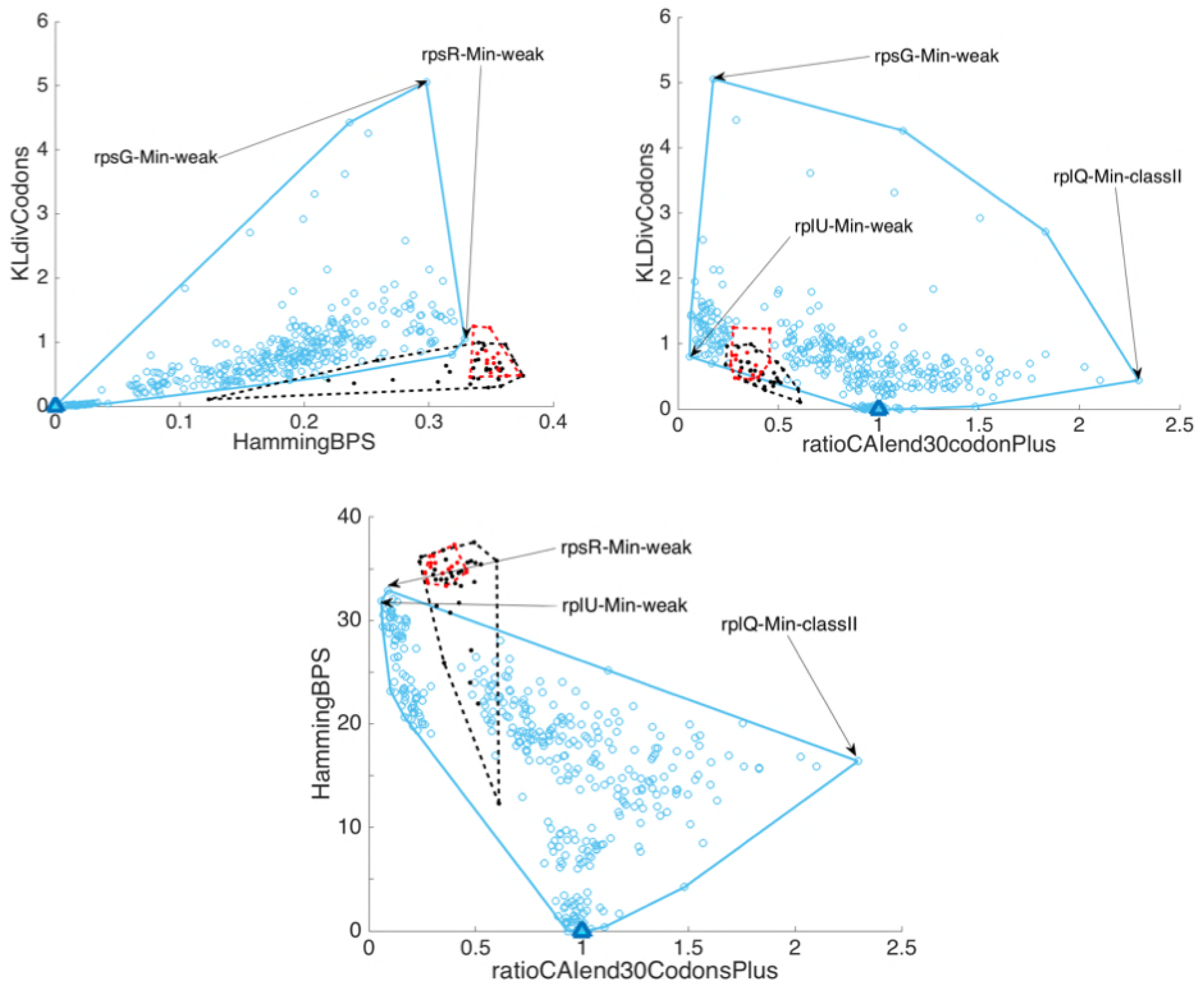
Pairwise convex hull plots (blue) in the three feature dimensions, selected in both the GC and SVM feature selection routines, circumscribing the entire data set of 360 modifications of 40 genes (i.e. modified with the nine, novel algorithms). Features from rplU, rpsG, and rpsR, modified with the Min_weak_ algorithm, and rplQ, modified with the Min-classII algorithm, lie at the corners of the convex hull bounding the data that requires predictions. *In vivo* experiments with *E. coli* to determine whether these genes with these changes impactimpact colony formation would provide new training data for the classifier that would span the feature domain more comprehensively and decrease extrapolation error for predictions made on this data set.

The algorithms span a spectrum of extremity from Sub-11 in which only 11 codons are eliminated, to tRNA_sub1_, in which 28 codons are eliminated, to Min-GCmax and Min_weak_, in which 43 codons are eliminated. The extremity of the codon changes induced by the algorithms is reflected most in the convex hull’s distance from the wild-type origin. As with Min_weak_ and tRNA_sub2_, the extreme algorithms that are not predicted fail also have convex hulls similar to each other in the feature space. Because they all eliminate more than half of the available pool of codons, their convex hulls are farther away from the wild-type origin than the less extreme algorithms. These extreme codes have, on average, a lower KLdivCodons and a higher ratioCAIend30CodonsPlus, compared to the codes that are predicted to result in design failure. It is also interesting that one gene, rpsG, disproportionately impacts the shape of the convex hull plots for all algorithms when KLdivCodons is plotted (i.e. it is an outlier). The KLdivCodons of rpsG is much higher for each algorithm than any other modified gene. While changes to rpsG from the nine codes are not predicted to fail, according to our classifier, it is important to note that the area of the feature space occupied by these modifications of rpsG is distant from that occupied by the training set, implying increased likelihood of extrapolation error for these predictions. Better predictions for these modifications of rpsG might be obtained if the classifier’s training set is expanded to include colony formation data in the area of the feature space its modifications occupy.

Conducting *in vivo* experiments to collect more training data is not a trivial endeavor, however. The *in vivo* experiments in [5] which produced the data used to train the classifiers in this work, though less difficult and expensive than full, genetic-code engineering *in vivo* experiments, were challenging and time consuming. In this context, discovering a minimal set of “high-value,” new examples that can be used to improve the model would be advantageous. Figures 10 a-c. are pairwise convex hull plots (blue) in the three feature dimensions, selected in both the GC and SVM feature selection routines, circumscribing the entire data set of 360 modifications of 40 genes (i.e. modified with the nine, novel algorithms). Features from rplU, rpsG, and rpsR, modified with the Min_weak_ algorithm, and rplQ, modified with the Min-classII algorithm, lie at the corners of the convex hull bounding the data that requires predictions. *In vivo* experiments with *E. coli* to determine whether these genes with these changes impactimpact colony formation would provide new training data for the classifier that would span the feature domain more comprehensively and decrease extrapolation error for predictions made on this data set.

These convex hull visualizations intuitively determine and display minimal sets of informative data that expand the domain of the feature space covered by the training data. Other more commonly used metrics for determining valuable data to collect to improve classifier performance, such as uncertainty sampling and expected error reduction [32], could also be used in this visualization to improve researchers’ intuition about where valuable data defined by this metric lives in the feature space. In principal, this technique can be used to display good training data for any classifier and any set of genes modified under any set of algorithms.

In general, all of these visualizations can be used to gain a synoptic view of the impact of modifying codon-usage with different algorithms in large groups of genes. In particular, these plots can be used to answer the following questions: How do algorithms differ relative to each other in terms of size, shape, position of and data spread within their convex hulls? How different, on average for a large set of genes of interest, is a sequence modified with a particular algorithm from the wild-type sequence? Is the algorithm and set of genes we are interested in gauging close to or far away from the data used to train the design success/failure classification model (i.e. how good will model predictions be for a given algorithm implemented on a given set of genes)? What are the most valuable experiments to conduct to obtain training data that will improve model predictions? Answers to these questions augment and complement the information obtained in a binary classification based on codon modifications in a single gene. These plots can thus be used in tandem with the classification models to flesh-out the landscape of how different genetic code algorithms influence large numbers of salient genes in the genome.

## Discussion

With a small dataset of 47 experimental examples, we have developed classifier models that estimate which re-coding algorithms, applied to a highly expressed, essential genes, will result in a design failure. These results are significantly better than random guessing at the 5% level. Furthermore, the AUC values were determined using a nested cross validation approach to maximize generalizability to unseen data. Given the extensive resources required for *in vivo* recoding experiments, we will now use results from this simple model to help eliminate problematic experimental trajectories.

We anticipate that this model will improve as more *in vivo* data is collected for training. Future work will involve iteratively conducting promising experiments, based on model predictions, and using this data to train better models for predicting the next round of experiments. In particular, our training set was biased by the fact that only the FCS and FC algorithms were used on a set of 40 highly expressed, essential genes. In addition, data from different algorithms needs to be collected and used to make a more robust model. While data from the genes used here are likely to prove useful in determining whether a recoding algorithm is problematic, other highly expressed and/or essential genes should also be tested. Feature engineering efforts to develope new, promising features is expected to improve results as well.

Our classification methods predict effects of modifying codon usage on a gene-by-gene basis. It is possible that characteristics of the convex hulls plotted in Figures 9. and 10. could yield insights into how modifications in multiple genes *in vivo* would combine to impactimpact viability. To verify this, we will need *in vivo* data in which such multi-gene modifications have been engineered. Such experiments are ongoing in other research groups [6]. Finally, the other dimensions in the multi-objective optimization promised by genetic code engineering remain to be explored. How different does a recoded organism have to be to confer genetic isolation and viral resistance? What are the bounds on incorporating non-natural amino acids into an organism’s protein repertoire? And, can the principles of recoding be successful in more complex organisms with much larger genomes?

## Materials and Methods

### Model Testing and Training Data

The modeling and performance evaluation methodology when building predictive models differ from that involved in constructing explanatory/causal models [14]. We make no claims in this paper about causality or identifying mechanisms. The scope of the modeling in this work is focused on the pragmatic goal of building an *in silico* pipeline to help predict design success or failure, as a binary yes/no variable, when the codon usage in single genes is changed according to a specific recoding algorithm. In this sense, the tools we present here are decision-support tools for genetic code engineers. The intention of this work is not to build upon or infer biological mechanisms-it is to facilitate and inform the scientific experimentation process in genetic code engineering. This specific goal informed the types of features we examined, the validation methodology we used to estimate the performance of our model and our approach to correlated variables in the model.

When choosing the features we tested for predictive power, our focus was on whether the features were measurable and had predictive potential, rather than on whether such features described possible causal mechanisms derived from theory. Additionally, our validation metrics were empirically derived through nested cross-validation, instead of theoretically derived metrics of goodness-of-fit that are used for regression models when the magnitude and confidence of feature weights in the model are used to infer causal relationships. Finally, correlated predictors do not present the same problems in prediction as they do in explanatory modeling, where understanding the causal strength of a predictor is impacted if other variables being inspected are correlated with it [14,15]. As a consequence, we did not preclude correlated features in our model; rather, we examined whether their presence had a consistent, significant effect on classifier performance.

For feature selection, model training and cross-validation, we used a dataset consisting of sequence data from altered genes, tested *in vivo*, and the associated observations of colony formation (e.g. failed or succeeded), from the experiments as described in [5]. 16 of 42 modified genes implemented *in vivo* in the experiments in [5] resulted in no observed colonies. As a result, a more conservative codon-usage algorithm was tested for these genes in which the 51 allowed codons remain unchanged in the coding sequence. Furthermore, any synonym that enabled successful colony formation from the allowed, synonymous codons was used to replace forbidden codons when sampled synonyms resulted in failed colony formation [5]. This codon-usage algorithm is referred to as the “The Forbidden Codon” (FC) algorithm in this work. All *in vivo* implementations of FC algorithm were viable.

Information was not available to us for all modified genes in [5]; therefore, data in our *in silico* work consists of a subset of the genes (40 out of 42) described in [5]. In this subset, forty of the modified genes were changed according to FCS algorithm in Figure 2. These genes are listed in Table 1. in regular type (e.g. not bolded). Of these, sixteen failed to produce colonies after *in vivo* implementation in [3], cf. **Design Failure** row in Table 1. Twenty-four resulted in successful colony formation, cf. **Design Success** row, regular type in Table 1. Seven of the sixteen genes in our dataset that were changed with the FCS algorithm and resulted in failure, were modified instead with the more conservative “Forbidden Codon” (FC) algorithm, in which only forbidden codons are replaced in subsets to test for viability, and implemented *in vivo* [3]. Modifications in this set of seven genes resulted in successful colony formation under the FC algorithm but failed colony formation under the FCS algorithm, which is why they appear in both the **Design Failure** and **Design Success** rows in Table 1.

To minimize potential sources of extrapolation error, we modified, *in silico*, the same set of forty genes with our nine, novel codon-usage algorithms. The sequence data for wild-type sequences that we used for our features was obtained from EcoCyc, an *E. coli* Database [16]. Because interactions between the initial protein coding sequence and the ribosomal binding site (RBS) in genes with modified codon usage could potentially influence viability, we included 19 base pairs prior to each start codon. We appended these sequences to the wild-type and codon-modified protein-coding sequences of all genes to compute features using these extended sequences.

### Feature Selection, Model Learning and Validation Methodology

We computed twenty-two features for each wild-type and codon-modified (e.g. by the FC, FCS, and the nine novel codes) sequence. We chose several features quantifying codon bias and mRNA folding free energy based on observed physiochemical consequences and proposed mechanisms identifying the role of synonymous codon changes in translation [13,17-22]. These features were further partitioned and computed for salient subsets of codons in the genes. For example, several studies identify the first 30-60 base pairs of the coding sequence as a slowly-translated “codon ramp” relative to the remainder of the gene [13,17-22]. Based on this information, we computed codon adaptation indices (CAI) for the first 30 codons of each gene and all subsequent codons prior to the stop codon, each as independent features. Since the estimated interval of codons included in this ramp is not precise and there are possibly other salient codon subsets within the gene, we also computed CAI for the 21^st^ codon (including start) to the 21^st^ codon before the stop codon and for the 13^th^ codon (including start) to the codon before the stop codon. We choose these intervals to explore the influences of groups of codons approximately in the middle and at the end of the genes. A sliding window approach was leveraged in [23] to explore gene subsets for these types of features more comprehensively. To keep our feature space small, given our small dataset, we did not use the same approach here. Future studies with more data will be able to accommodate similar, more comprehensive feature spaces. Several reports, [3,22-24], discuss the influences of mRNA structure around the ribosome binding site and the beginning of the codon sequence on gene-expression levels. Based on these results, we also computed mRNA folding free energy for the coding sequence alone, the coding sequence plus the RBS and the RBS plus the first ten codons in the coding sequence.

Other features were theoretical metrics of information changes in the modified genes, where information is measured using alphabets of codons or base pairs, relative to the wild-type gene sequence. Feature abbreviations and descriptions are given in Table 2. The KL divergence computations on codon histograms from each gene were computed using additive (Laplace) smoothing [25-27] to address the fact that not all possible codons were observed in every gene. All features, other than those comparing codon and base pair frequencies in the genes, were computed as a ratio of the feature from the modified sequence divided by the feature computed for the wild-type sequence. This was done to assure all features spanned a similar range. Dividing by the length of the gene normalized the Hamming distance and we used a normalized mutual information metric that bounds the values within [0 1]. KL divergence values remained within [0 3] for all genes changed with all codon-usage algorithms. Normalization was unnecessary for this feature.

**Table 2.** Feature descriptions. ^*^ CAI features are computed for subsets of the gene informed by observations of a “codon-ramp” about 30-60 base pairs into the coding sequence [13,17-22] and by possibility of other salient subsequences. ^**^ mRNA folding free energy features are computed for subsets of the genes informed by observations of the influence of secondary structure between the RBS and different parts of the coding sequence on expression levels and translation rates [22-24].

| Information Theoretic/Statistical Features |  |
| --- | --- |
| name | description |
| normMIBPS | normalized mutual information between the base pair counts in the wild-type coding sequence and the modified coding sequence interpreted where the base pair names are interpreted as four exchangeable cluster labels |
| normMICodons | normalized mutual information between the codon counts in the wild-type coding sequence and the modified coding sequence interpreted where the codon names are interpreted as 61 exchangeable cluster labels |
| entBPS | entropy of the base pair counts in the coding sequence |
| entCodons | entropy of the codon counts in the coding sequence |
| KLdivBPS | KL-divergence between the wild-type and modified base pair distribution of base pairs in the coding sequence |
| KLdivCodons | KL-divergence between the wild-type and modified codon distribution of base pairs in the coding sequence |
| HammingBPS | Hamming distance between the wild-type and modified base pair distribution of base pairs in the coding sequence |
| HammingCodon | Hamming distance between the wild-type and modified codon distribution of base pairs in the coding sequence |
| countA | count of adenine in protein coding sequence |
| countC | count of cytosine in protein coding sequence |
| countG | count of guanine in protein coding sequence |
| countU | count of uracil in protein coding sequence |
| countAplusU | count of uracil and adenine in protein coding sequence |
| countGplusC | count of guanine and cytosine in protein coding sequence |
| CAI | codon adaptation index (CAI) for the entire gene |
| CAIbeg* | CAI for codons in the first 30 codons in the gene (excluding the start codon) |
| CAIend30codonsPlus* | CAI for codons from the 31<sup>st</sup> codon to the final codon (before the stop codon) |
| CAImid* | CAI for codons from the 21<sup>st</sup> codon (including start) to the codon 21 codons before the final codon (before the stop codon) |
| CAIend* | CAI for the 13<sup>th</sup> codon to the last (before the stop codon) |

| Physiochemical Features |  |
| --- | --- |
| name | description |
| RNA | mRNA folding free energy for the protein coding sequence |
| RNARBS** | mRNA folding free energy for the protein coding sequence and the ribosomal binding site |
| RNAFoldRBS30** | 19 base pairs before the start codon plus the first 30 base pairs of the coding sequence |

To maximize generalizability of our performance estimates, we implemented a nested cross-validation (CV) involving two tiers (cf. Figure 2 and Figure 3). In the outer tier, depicted in blue in both figures, a test set of ten was selected randomly without replacement from an exhaustive set of all unique combinations of ten observations including five sequence changes associated with a design failure and five sequence changes associated with a design success. This left thirty-seven observations for training the model in the outer tier and for use in an inner CV tier, green and gray in both figures, to learn optimal parameters for the model (meta-learning). The inner tier consisted of exhaustive leave-pair-out cross validation (LPOCV), holding out one observation with a design success and one with a design failure, on each iteration. This CV tier was used to optimize three meta-parameters: the number of features {4,5,6,7} to be included in the model, the threshold on the soft classification for binary design success/failure prediction and whether to include correlated features in the selected set or not.

To eliminate correlated features when testing whether their presence impactimpacts classifier performance, if two feature had a Spearman’s correlation higher than |0.6|, we removed the feature with the smallest Cohen’s d value from the set of features used in the feature selection process. Sets of four to seven features with the largest magnitude Cohen’s d effect size in design success/failure discrimination were selected for each inner CV training set of 35 and the test pair was classified using selected features. The number of features with the best area under the ROC curve (AUC) from the LPOCV was used as the number of features to be selected in the outer CV step using the training set of 37.

Finally, from the ROC curve produced from the inner CV loop with the best AUC, we also identified a classification threshold. We chose this threshold to occur at the operating point at which we reach 90% sensitivity with the fewest possible false alarms. We chose this point because this tool is intended to decrease the risk of running genetic code experiments with codon-usage modifications likely to result in design failure. Misclassifying a significant number of codes that would result design success as resulting in design failure does not entail as much risk in terms of lost time and money. In other words, type II errors are costlier than type I errors in this domain and the classification threshold was set to account for the asymmetric risk.

A Gaussian classifier and a soft-margin SVM with a linear kernel were trained using these data and these features. We chose simple classifiers because our dataset was small and complex classifiers are more likely to over-fit to spurious patterns in the training data. These models were then used to predict the classes of the ten held-out data points in the outer CV loop. This loop was iterated 1000 times for 1000 different subsets of 10 held-out observations. AUC, sensitivity and specificity was computed for each of the 1000 iterations and averages over each metric estimate overall performance of the SVM and GC on this dataset. The complexity and rigor of this validation methodology was necessary to achieve generalizable performance estimates with our small dataset.

### Definition of Nine Novel Codon-Usage Algorithms

As mentioned above, we used our model to predict *E. coli* viability when the set of forty genes from Table 1. were modified with nine novel codon-usage algorithms. All of the algorithms presented here are *reductionist* algorithms, i.e. some codon assignments to amino acids have been removed, leaving those codons blank—unassigned to any amino acid. In contrast, a *reconfigured* algorithm would change the assignment from one amino acid to another. We consider reductionist algorithms an important step toward implementing more complex reconfigured algorithms in living cells.

Previous efforts [1,2]_reported the removal of one codon throughout the entire genome of MG1655, called the r*E. coli* algorithms. The algorithms designed and evaluated here sample greater extremes, from an intermediate number of changes (removing 11 codons) to bare minimum codes (leaving only one codon for each amino acid, removing roughly two-thirds of the natural 64 codons).

#### Sub_11_

This set subtracts 11 codons from the canonical genetic code. These codons are used at low frequencies within MG1655 genes. We hypothesize that this set would remove enough codons to provide benefits such as strong resistance to bacteriophage infection, while still providing enough choices of synonymous codons to retain flexibility for troubleshooting emergent challenges with genome design and implementation. Modification of the codon usage in the genome combined with deletion of the corresponding tRNA genes is expected to be sufficient for the genomic installation of this code.

#### tRNA_sub1_

One version of a code that seeks to minimize the number of naturally-occurring tRNAs retained in the design. With 28 forbidden codons (i.e. 36 natural assignments retained) each amino acid is encoded by one or two codons.

#### tRNA_sub2_

A stricter version of a code that minimizes the number of naturally-occurring tRNAs required to represent each amino acid. With 36 forbidden codons (i.e. 28 natural assignments retained) each amino acid is encoded by one or two codons (more often one codon, compared to <u>tRNA_sub1_</u>).

#### Min_curated_

The first of several designed codes that theoretically only use one codon to represent each of the standard 20 amino acids. In all such minimal codes (“mincodes”) 43 codons are forbidden (21 codons remaining, i.e. for natural amino acids plus one stop codon). The mincodes are thus deterministic in the sense that once a gene product is defined at the amino acid sequence, the DNA sequence for the gene is definitively known. Leaving the genome engineer with no choices of DNA sequence flexibility is a considerable drawback for any eventual troubleshooting processes. In this report, all mincodes described are subtractive, only removing codon assignments, not reassigning or re-organizing. The primary means to implement this code would be to extensively modify codon usage in the genome, followed by deleting a large proportion of tRNA genes to be used no longer. However, in order to cleanly remove all forbidden codons from the genetic code, the remaining tRNA genes would also have to be significantly engineered to recognize exactly one codon each (instead of two or three each, which is most typical). This first mincode, <u>Min_curated_</u>, is labeled “curated” to denote that several different design principles were considered and integrated by the designer. First, the most commonly used codons in MG1655 were emphasized, with the intent of not disrupting translation of highly expressed essential genes. Several decisions were also made for the purpose of making this code more facile to implement using synthetic DNA. To minimize runs of homopolymers that can be more challenging for DNA synthesis and high throughput sequencing, codons TTT, CCC, and GGG were forbidden. Pairs of codons that could give rise to palindromic repeats were also minimized, to reduce unintended RNA secondary structures as well as to avoid mispriming events during *de novo* DNA synthesis.

#### Min_classII_

The 21 codons retained in this mincode are those defined as most utilized in highly expressed *E. coli* genes, as categorized by Hénaut and Danchin (“Class II”) [28].

#### Min_sparse_

43 codons are forbidden, with the remaining allowed codon assignments chosen specifically to maximize the genetic distance between codons. Put another way, most single base mutations give rise to a forbidden codon. For example with the allowed alanine codon (GCG) only one of the possible nine single mutations gives rise to another allowed codon (to CCG, proline). Thus, the majority of spontaneous single-base mutations that arise would be expect to yield a null (i.e. unassigned) codon, disrupting translation and giving rise to defective protein products.

#### Min_ATmax_

The 21 codons of this mincode were chosen in order to yield the highest possible content of A:T base pairs. This design principle was chosen in order to explore one extreme of thermodynamics (e.g. self-folding energies)

#### Min_GCmax_

The 21 codons of this mincode were chosen in order to yield the highest possible content of G:C base pairs. This design principle was chosen in order to explore the other extreme of thermodynamics (e.g. self-folding energies)

#### Min_weak_

The 21 codons of this mincode were chosen according to the codons least used in MG1655 for each amino acid. Thus genes modified according to Min_weak_ are expected to require the most extensive degree of change from the wild-type sequence.

To be fully implemented in living cells, all minimal algorithms would also require extensive engineering not only of genome-wide codon usage in protein-coding reading frames, but also of the tRNAs that recognize these codons. Otherwise many forbidden codons would still be recognizable by the remaining tRNAs due to wobble base-pairing between codons and anticodons. See Table 3. for a synoptic depiction of these nine codes as well as the FC, FCS and *rE. coli* codes.

**Table 3.**
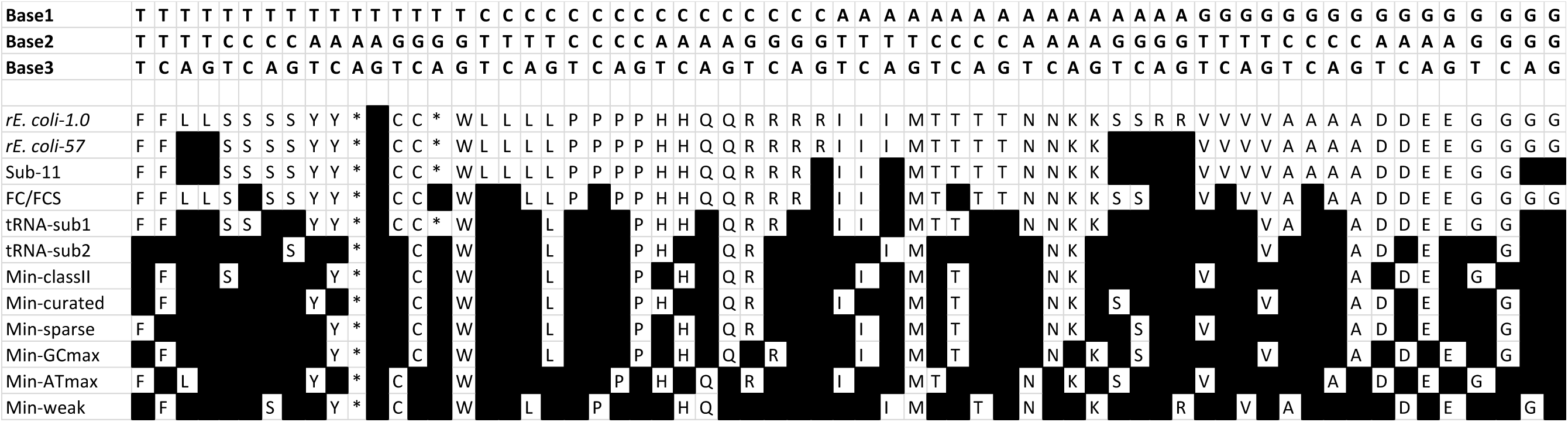
Depiction of the nine, novel codes as well as the *rE. coli*-1.0, *rE. coli*-57, FC and FCS codes. Blacked out boxes indicate disallowed codons for a given amino acid and codes are shown in descending order, from the fewest to the most codons removed. Standard symbols for amino acids are used.

**Table 4.** Performance metrics along with 95% confidence intervals for the SVM and GC classifiers. These results demonstrated that there was information in the selected features to predict design success/failure due to codon-usage modifications from the FCS and FC algorithms. Furthermore, at the 5% significance level, the GC AUC was better than that of the SVM with a linear kernel.

| Classifier | Mean Sensitivity<br>[95% CI] | Mean Specificity<br>[95% CI] | Mean AUC<br>[95% CI] |
| --- | --- | --- | --- |
| SVM | 0.76<br>[0.48 0.93] | 0.52<br>[0.33 0.70] | 0.68<br>[0.51 0.85] |
| GC | 0.80<br>[0.54 0.95] | 0.48<br>[0.30 0.67] | 0.72<br>[0.55 0.88] |

**Table 5.**
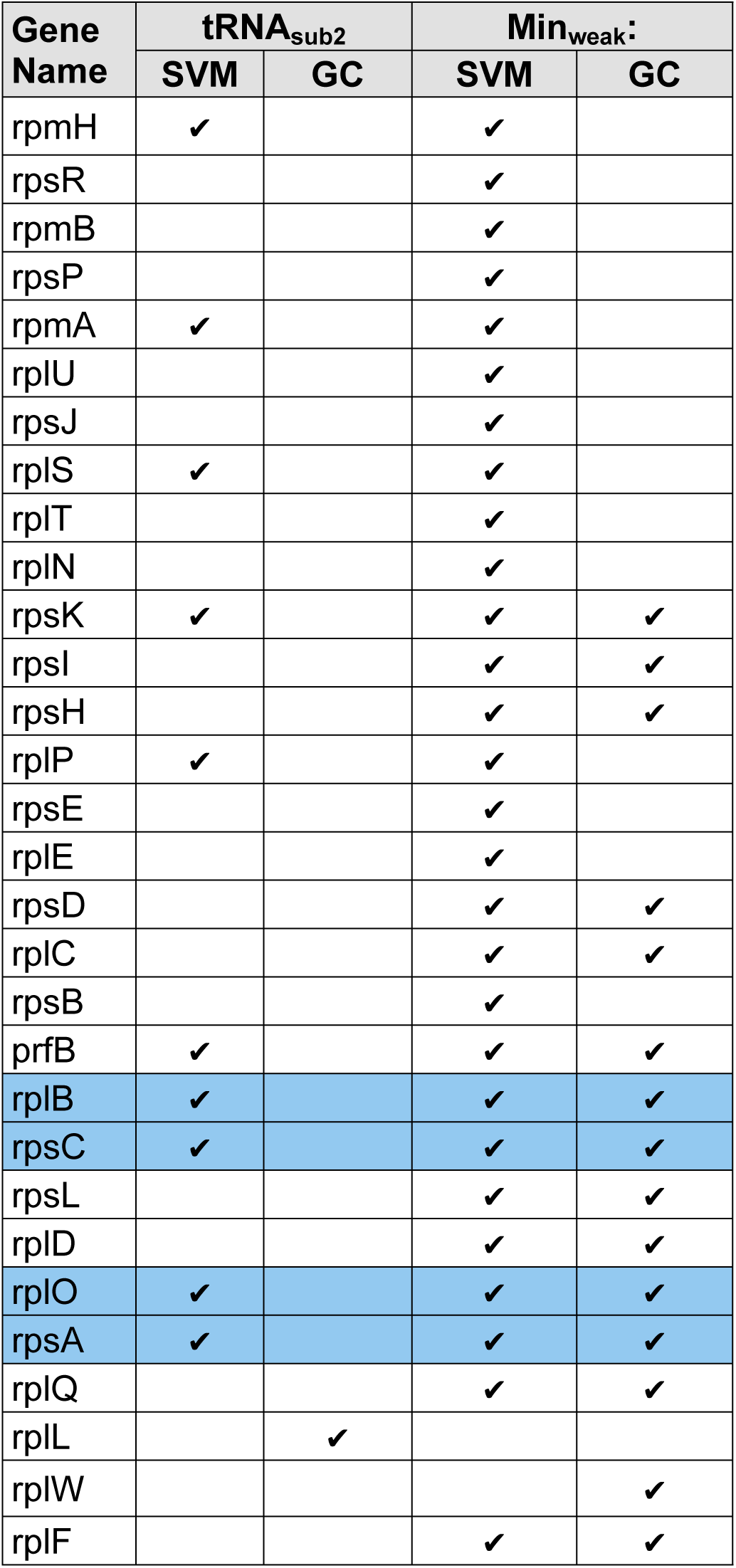
Genes modified with particular codes that were classified as design failures using our model trained on the 40 genes modified with the FC and FCS algorithms. Of the nine algorithms we tested, only two are predicted fail, tRNA_sub2_ and Min_weak_. Min_weak_ was the most disruptive to colony formation, according to our model, since many genes modified with this algorithm were classified as failing by both the GC and SVM. Several genes (highlighted in blue) appear to be more sensitive being modified than others. These genes are predicted to result in a design failure under both tRNA_sub2_ and Min_weak_ for the SVM and are classified failing by the GC under Min_weak_. They also resulted in a design failure when modified with the FCS algorithm and implemented *in vivo* in [3]. These may be good “bellwether” gene candidates in the sense that different modifications to them appear more likely to fail.

Next, we learned a model trained on the entire FC and FCS dataset to be used to classify the new data from the nine codes described above. We determined which features were most predictive design suc
cess/failure classification and trained SVM and GC classifiers. Model parameters, including the number of features to used for classification, whether to include correlated predictors or not and the classification threshold, were learned using a LPOCV, as in the nested cross-validation study described above.

### Software/Hardware

We performed *in silico* codon usage modifications for the nine novel codes and most feature computations with scripts written in Python and packages from Scikit Learn (normalized_mutual_info_score) and Scipy (stats). To compute the mRNA folding free energy features, we used an open-source package called The Vienna RNA Package [29]. To efficiently implement the expensive nested cross validation pipeline for the SVM and GC, we used Matlab code and pMatlab [30], a capability enabling Matlab parallel processing across hundreds of cores, and the MIT/MIT Lincoln Lab supercomputing facilities at Holyoke, MA.

## Acknowledgements

All experimental data we used to test and train the classifier was collected during the experiments reported in [5] and shared with us by Dr. Marc Lajoie. We are extremely grateful for his contributions to this work. These also include extensive and very insightful feedback on the manuscript and results. We also thank Dr. Nicholas Guido for his detailed review of the work and helpful comments, Anna Shcherbina for providing a Python code template to use for substituting codons *in silico* and Siddharth Samsi for providing a code template for running the nested cross validation in parallel using pMatlab [30] on the MIT Lincoln Laboratory supercomputing cluster. This work is sponsored by the Assistant Secretary of Defense for Research & Engineering under Air Force Contract no. FA8721-05-C-0002. Opinions, interpretations, conclusions and recommendations are those of the author and are not necessarily endorsed by the United States Government. Research reported in this publication was supported by the National Cancer Institute of the National Institutes of Health under award number R01CA173712.

## Appendix

**Figure S1.**
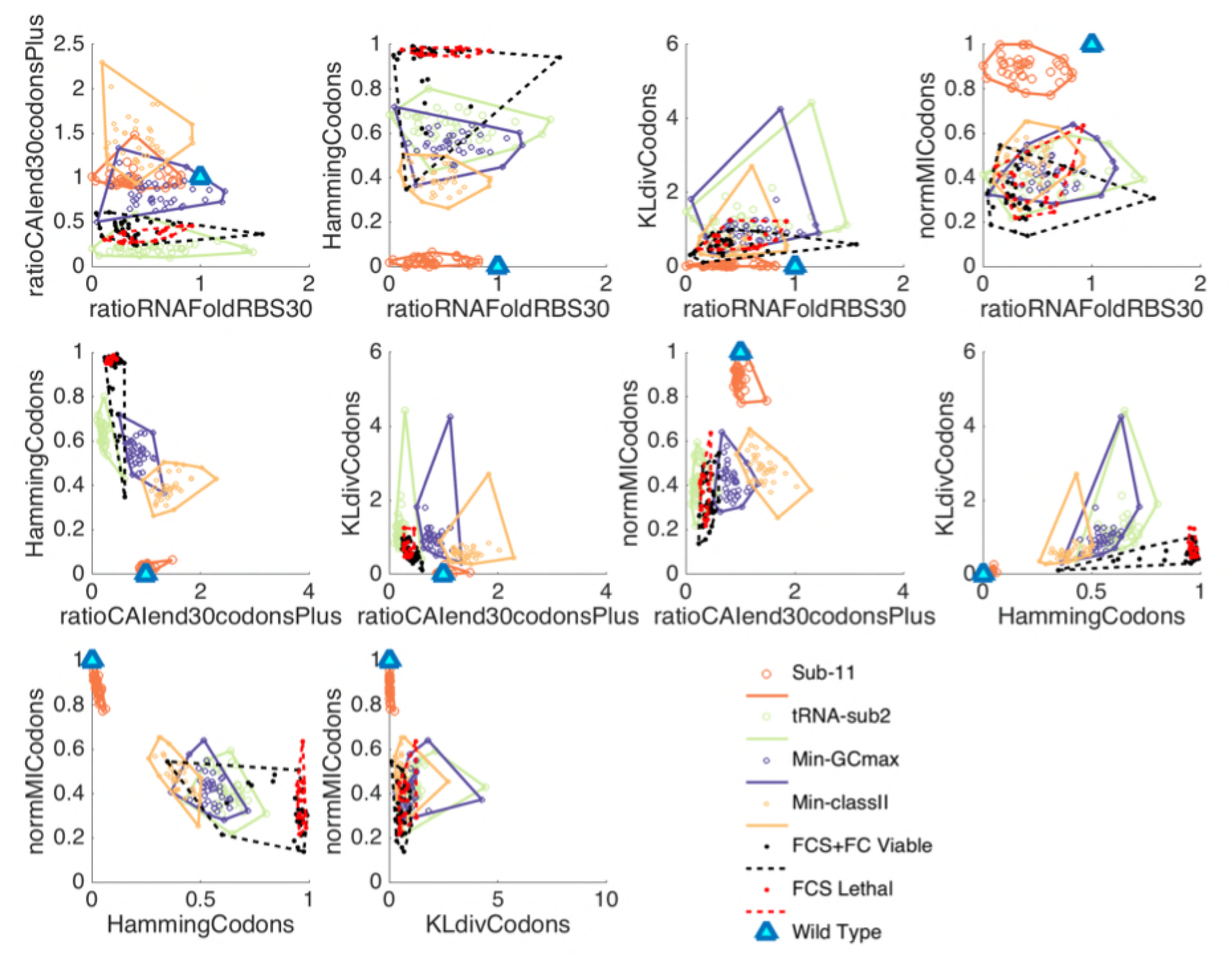
Convex hull plots for four of the nine novel codes we defined and investigated as possible candidates for genetic code engineering. All five features selected in the uncorrelated feature selection routine for the SVM are plotted pairwise. Note that the convex hulls of the genes recoded with the tRNA-sub2 algorithm, one of the two algorithms predicted to result in a design failure by the two classifiers, are consistently closest to the lethal training examples. Min-GCMax and Min-classII convex hulls overlap significantly in these plots, indicating that the two algorithms effect the forty genes similarly in the feature space salient for design failure prediction. The Sub-11 convex hull is separate from those of the other codes and closest to the wild type origin. It is also the smallest convex hull in all feature plots, signifying that all forty genes are similarly influenced by this code.

**Figure S2.**
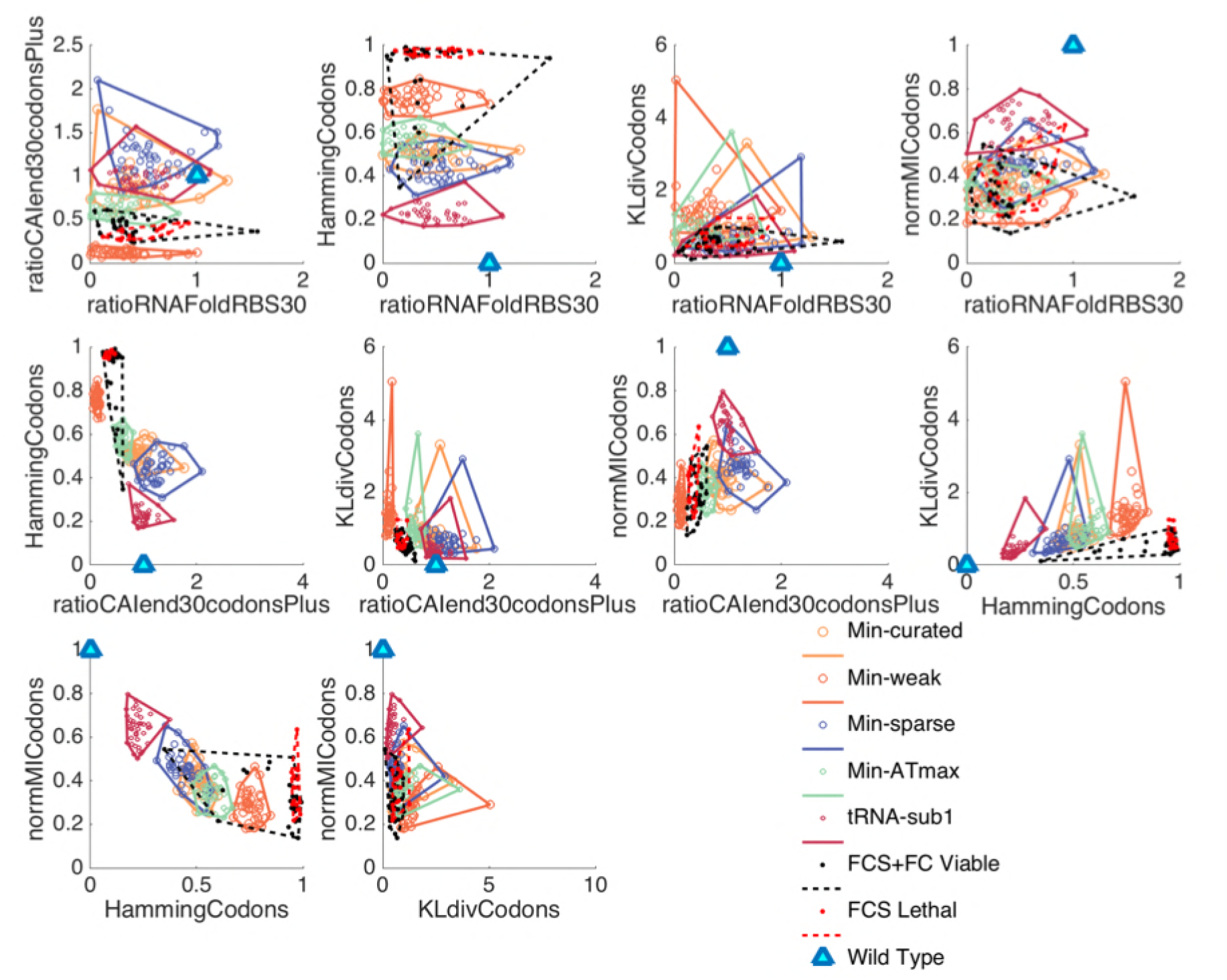
Convex hull plots for the five of the nine novel codes not depicted in Figure S1. The Min-weak convex hull is closest to the lethal training examples and is the smallest in most of the feature plots. This indicates that, when recoded with this code, the forty genes are effected similarly each other and to the lethal training examples that were recoded using the FCS algorithm. There is significant overlap among the Min-sparse and Min-curated convex hulls. These codes effected the forty genes similarly in the salient feature space, both in terms of the absolute placement and of the size and shape of the convex hulls. Min-ATmax effects the genes in a somewhat similar fashion but, on average, has a smaller convex hull in the pair wise plots, indicating that the genes were more consistently effected in the same way relative to wild-type with this code. tRNA-sub1, the code with the fewest disallowed codons, is always the closest to the wild-type origin, relative to the other convex hull plots.

